# The origin of mechanical advantage in angiosperms

**DOI:** 10.64898/2026.05.22.727130

**Authors:** Anju Manandhar, Scott A. M. McAdam, Fulton E. Rockwell, Yiling Fang, Timothy J. Brodribb, N. Michele Holbrook

## Abstract

- Mechanical interaction between guard cells and epidermal pavement cells enables large stomatal apertures and high productivity in angiosperms. We do not know when this response evolved, but over the last 169 years we have found that mechanical advantage has been tested in at least 230 species from 85 families. To date no data on this trait exists among angiosperms outside magnoliids, monocots and eudicots.
- To resolve the evolutionary origins of this critical stomatal response we tested for mechanical advantage across 14 additional species including the earliest diverging lineages of angiosperms.
- We find that mechanical advantage, while variable in magnitude, is present in all angiosperm species that have been measured, including *Amborella trichopoda* sister to all angiosperms.
- This response likely evolved once in flowering plants, in the common ancestor of this clade, remaining widespread across angiosperms today. We hypothesize that angiosperms could not have realized the full potential of physiological innovations in water transport without the evolution of this key trait that increased operational stomatal aperture.

## Introduction

Angiosperms have the highest rates of photosynthesis among land plants and dominate the terrestrial biosphere accounting for roughly 90% of extant plant species (Hernández-Hernández & Wiens, 2020). Physiological innovations of the angiosperms that facilitate high rates of gas exchange include a large diameter-low resistance xylem, high density-reticulate leaf venation, and small highly dense stomata. Often overlooked, among the physiological traits necessary for high rates of photosynthesis, is the aperture of an open stomate. Angiosperms are able to achieve large pore sizes because the guard cells open by displacing neighboring epidermal cells to achieve a maximized elliptical pore (Rockwell & Holbrook, 2017; Westbrook & McAdam, 2021). This interaction between guard cells and epidermal cells is known as mechanical advantage, reflecting the greater impact of epidermal cell turgor on stomatal aperture.

A consequence of mechanical interactions between guard cells and epidermal cells is that stomatal conductance is a function of epidermal cell turgor (Delwiche & Cooke, 1977; Edwards & Meidner, 1979). In the event of sudden dehydration epidermal turgor is lost allowing guard cells to push further apart, increasing pore size and gas exchange; the inverse occurs if epidermal turgor rapidly increases (Mcadam & Brodribb, 2012). These hydropassive movements are commonly called wrong-way responses (WWR). The first observations of changes in stomatal aperture by von Mohl(1856) in 1856 were WWRs. von Mohl saw that high epidermal turgor had a closing effect on stomatal aperture and that epidermal dehydration in sugar solutions had an opening effect; a similar experiment by Stålfelt (1929) in 1929 confirmed these observations. WWRs following leaf excision were some of the first reported responses of stomata measured in intact leaves in the 1800s using hygrometers, also called hygroscopes (Krutitzky, 1882; Darwin, 1898; Darwin & Pertz, 1911). Since this time, in angiosperms, WWRs following rapid changes in hydration have been observed using a wide range of methods including excising the whole plant, leaves, or branches in air or under water; increasing evaporation (Lange *et al*., 1971a); disconnecting the roots from the soil (Suissa *et al*., 2023); exposing either roots, cut leaves or isolated epidermis to high osmotic solutions (Raschke, 1970a; Glinka, 1971; Naidoo & Willert, 1994); damaging neighboring epidermal cells or even piercing the vascular tissue with a hot needle (Linsbauer, 1916).

The anatomical structure that allows for mechanical advantage in angiosperms remains unidentified. One study associates leaf vein architecture with the variation in the effect of mechanical advantage on stomatal dynamics (Buckley *et al*., 2011). Recently, large epidermal cell sizes were found to be associated with greater mechanical advantage in non-monocot angiosperms (Pichaco *et al*., 2024). Despite this recent work the impressive diversity of stomatal and epidermal anatomy in angiosperms (Rudall & Knowles, 2013; Rudall, 2023) remains poorly associated with stomatal function or the presence of mechanical advantage(Buckley *et al*., 2011). Yet, several epidermal features might be associated with stomatal function. For example, the degree of epidermal wall sinuosity and cell size affect cell wall rigidity (Bidhendi *et al*., 2019) which could influence guard-epidermal interactions. Another feature that could impact guard cell movements is the number of epidermal wall junctions abutting guard cells specifically at regions where guard cells move laterally. A major exception to this limited understanding of how epidermal anatomy influences stomatal function are the stomata of the Poaceae in which mechanical advantage has been explicitly tied to guard cell and epidermal cell anatomy (Raschke & Fellows, 1971). Poaceae species have unique dumb-bell shaped guard cells associated with specialized subsidiary cells that exchange volume when stomata open and close, with an active decrease in subsidiary cell osmotic potential occurring as the pore opens (Raschke & Fellows, 1971; Franks & Farquhar, 2007). A similar paracytic subsidiary cell arrangement, where a pair of epidermal cells bracket guard cells (without epidermal wall junctions lateral to the guard cells) is commonly observed in the monocot relatives of Poaceae (Rudall *et al*., 2017) like *Tradescantia virginiana* (Commelinaceae) and other angiosperms. Unlike the Poaceae, paracytic epidermal cells in *T. virginiana* do not have a coupled increase in guard solutes and volume for stomatal opening (Anisiobi, 2020); but *T. virginiana* still has mechanical advantage and WWRs. Whether a paracytic arrangement and hence a lack of epidermal wall junctions abutting the lateral side of guard cells is important for mechanical advantage remains unknown.

Mechanical advantage is well-described across a diversity of eudicot and monocot species, but appears to be absent from gymnosperms (Brodribb & McAdam, 2013; McAdam & Brodribb, 2014) and most ferns and lycophytes, with the exception of species from the family Marsileaceae (Westbrook & McAdam, 2021). Mechanical advantage is hypothesized to have been selected for in Marsileaceae because it enables the highest photosynthetic rates among ferns at the cost of high minimum conductance and an aquatic ecology (Westbrook & McAdam, 2021). The origin of mechanical advantage in angiosperms remains unresolved. If mechanical advantage is associated with enabling high rates of leaf gas exchange, as suggested by the evolution of WWRs in Marsileaceae (Westbrook & McAdam, 2021), then it could be hypothesized that angiosperms from lineages characterized by low rates of leaf gas exchange, particularly the earliest diverging angiosperms (Feild & Brodribb, 2013), might not have mechanical advantage. Most of the early diverging ANITA grade of angiosperm species, sister to the eudicots and monocots, have low stomatal conductance and low productivity (Feild *et al*., 2003) with developmental and physiological traits that limit high photosynthetic rates (Feild *et al*., 2001). The ANITA grade species lack the high vein and stomatal densities associated with high assimilation(Brodribb & Feild, 2010; Feild *et al*., 2011). One hypothesis of angiosperm origins posits that the common ancestor of this group evolved in a low light, high disturbance and mesic environment, similar to the native environments of many extant ANITA grade species (Feild *et al*., 2004). Though many ANITA grade angiosperm species do not have the typical high productivity that accompanies mechanical advantage observed in eudicots and monocots, the presence of mechanical advantage could still be critical for stomatal opening responses in understory adapted species (Rockwell & Holbrook, 2017). There is increasing evidence showing mechanical advantage results in faster stomatal opening speeds under high VPD (Pichaco *et al*., 2024; Shapira *et al*., 2024).

Whether mechanical advantage occurs in ANITA grade angiosperm species is unknown. *Amborella trichopoda*, sister to all extant angiosperms, has ABA-mediated stomatal closure responses to increasing VPD (McAdam & Brodribb, 2015); ABA response to high VPD is considered a necessary right way stomatal closure mechanism to counteract a WWR in angiosperms. The presence of an ABA mediated stomatal closure response to high VPD in *A. trichopoda* suggests that this species might have mechanical advantage. We hypothesize that the ANITA grade angiosperms have mechanical advantage despite the recorded low rates of gas exchange. Here we conducted an extensive literature review to document all species in which a test for mechanical advantage has been performed. We supplemented this dataset with experimental investigation of mechanical advantage in *A. trichopoda*, four species of Austrobaileyales, and three magnoliid species. We used the presence of the WWR on leaf excision to infer the presence of mechanical advantage. We also determined whether epidermal arrangement relates to the presence/absence of mechanical advantage in ANITA grade angiosperms. For comparison with species anticipated to lack mechanical advantage we measured stomatal conductance following excision in a lycophyte and three gymnosperm species. We also included observations of stomatal responses to excision in six eudicot species including an early diverging eudicot species, *Persoonia falcata* (Proteaceae; Proteales); and a species with extreme epidermal cell sinuosity *Trochocarpa gunnii* (Ericaceae; Ericales) (Jordan *et al*., 2010).

## Materials and methods

### Literature survey

To determine the range of species in which mechanical advantage has been described we searched literature to find as many records of mechanical advantage as possible. The terms mechanical advantage or WWR have not been used universally through the history of stomatal studies that show mechanical advantage. In the absence of common terminology, we determined that a study demonstrated mechanical advantage if dehydration caused a transient stomatal opening, or an absence of this response if stomatal closure began immediately on dehydration. Rehydration causing instantaneous stomatal closure was also used as evidence of mechanical advantage. Studies showed stomatal opening or closing with various techniques including: microscopy observations of stomatal aperture, permeability of stomatal pores to liquids as a measure stomatal openness, rate of color change in cobalt paper to indicate transpiration rates, balance measured transpirational water loss, transpiration rates of stomatal conductance measured with hygroscope/hygrometer, porometer or gas analyzer (Table S1). The documented experiments to dehydrate plant tissue ranged from petiole/branch excision, removing plants with roots from the soil, replacing nitrogen gas with helox in a gas analyzer to increase leaf water loss, step increases in VPD experienced by the leaf, adding osmotic solutions to the soil or floating leaf tissues in osmotic solutions, damaging epidermal cells with a needle (Table 1). Experiments that hydrated plant tissue did so by excising petioles or branches in water, watering the soil, pressuring the soil and roots to increase water potential or floating leaf tissue in water (Table 1). We did not include data from the few early reports that exist in which increases in gravimetrically determined transpiration occurred in excised branches but the cut end was not sealed after excision; these increases in transpiration cannot be directly attributed to stomatal responses and are likely due to considerable water loss from excised stem ends as comprehensively discussed by Arland (1929). The Google translate application was used to translate German literature. The World Flora Online (wordlfloraonline.org) was used for modern taxonomic determinations, except in the case of the misnomer of *Senecio minimus* in the database and the debatable determination of *Pisum sativum*.

**Table 1.**
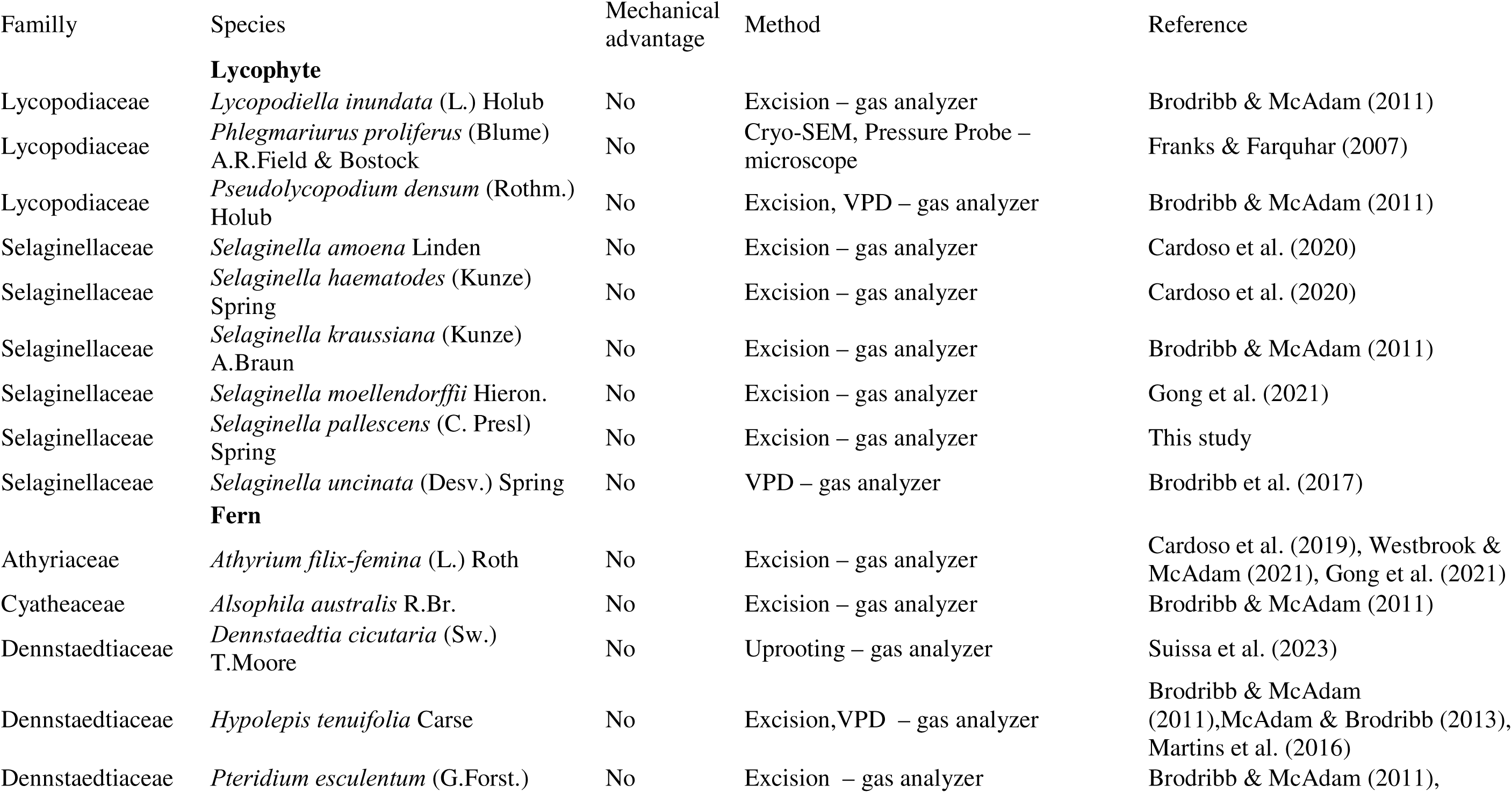
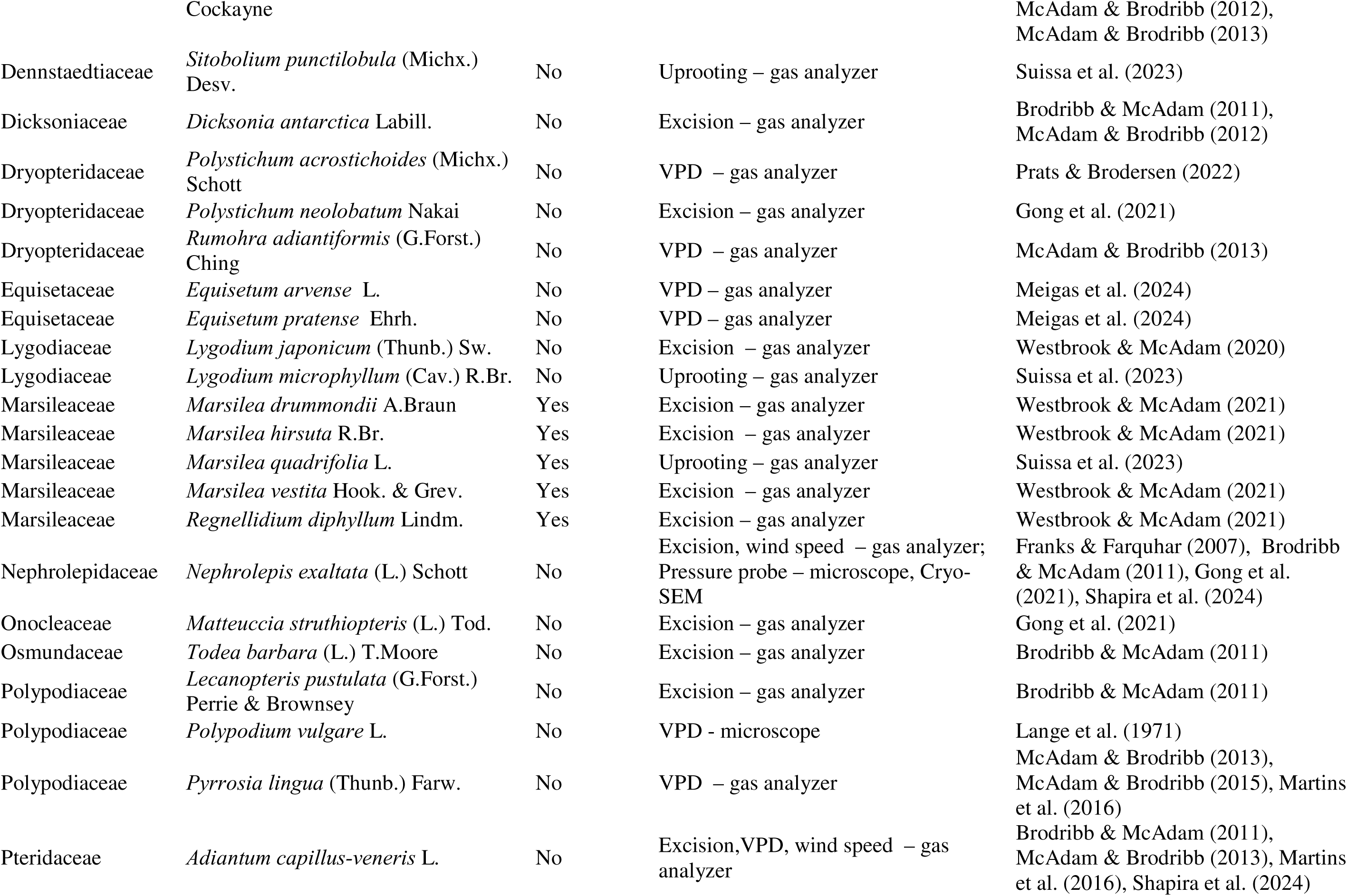
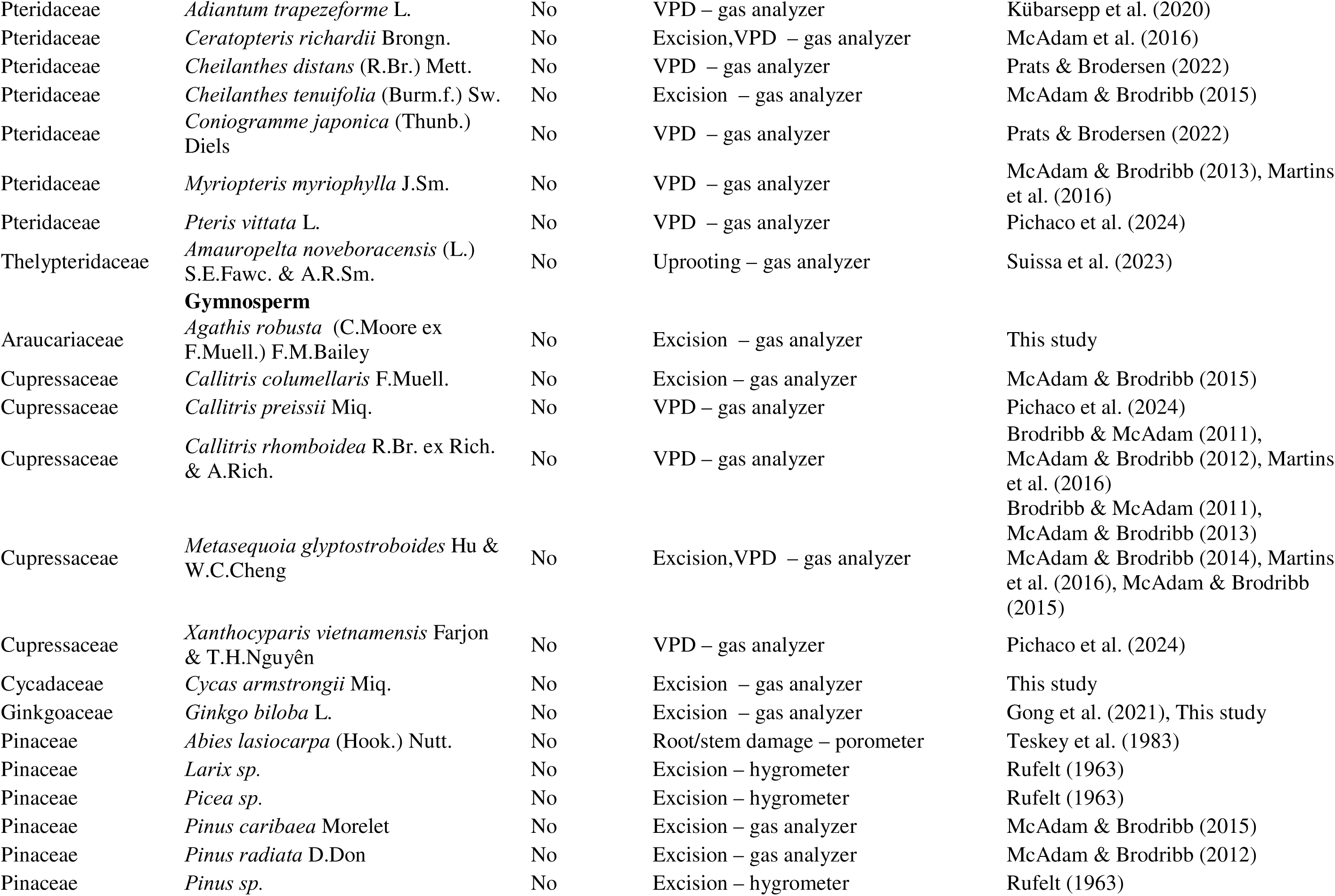
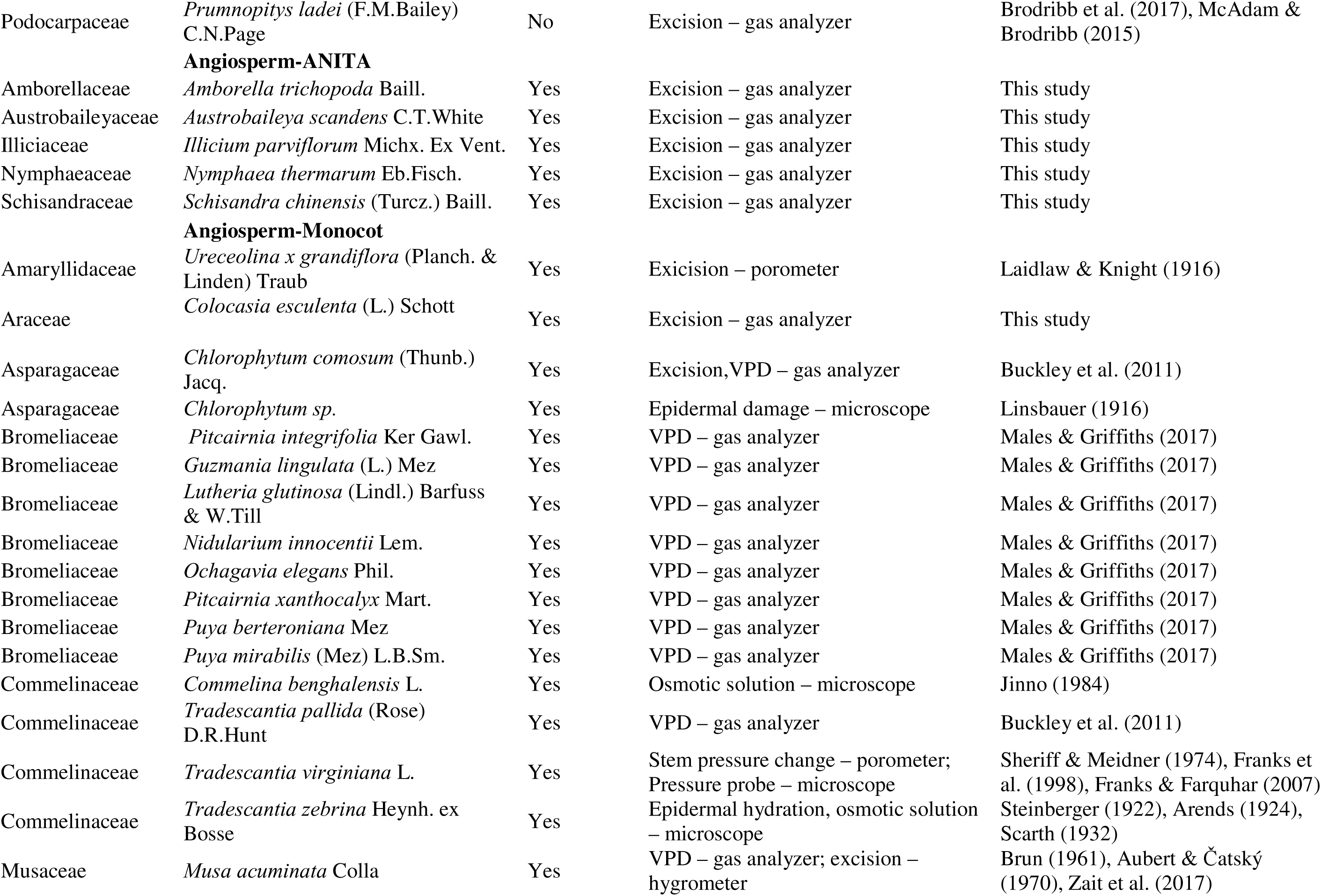
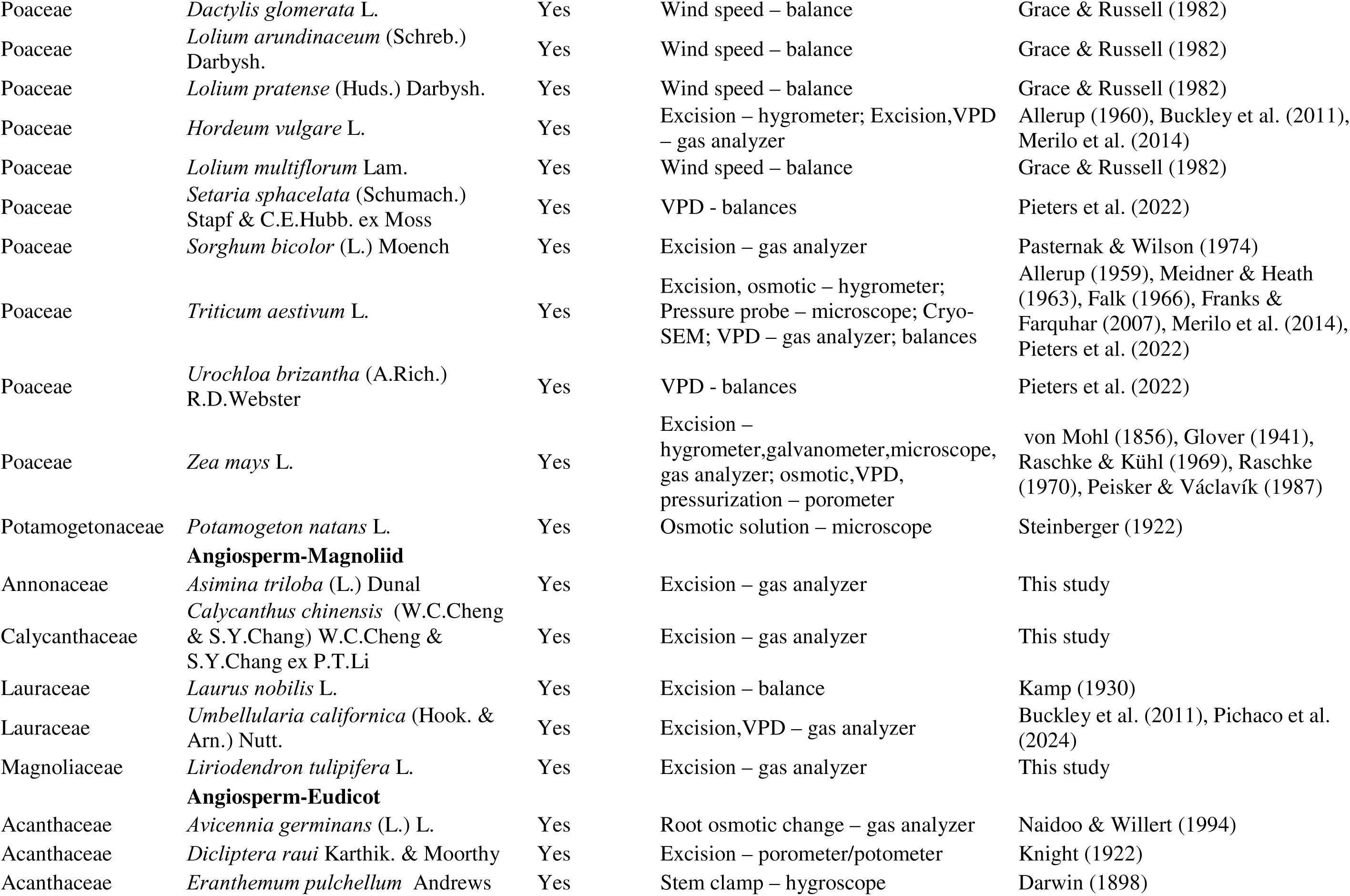
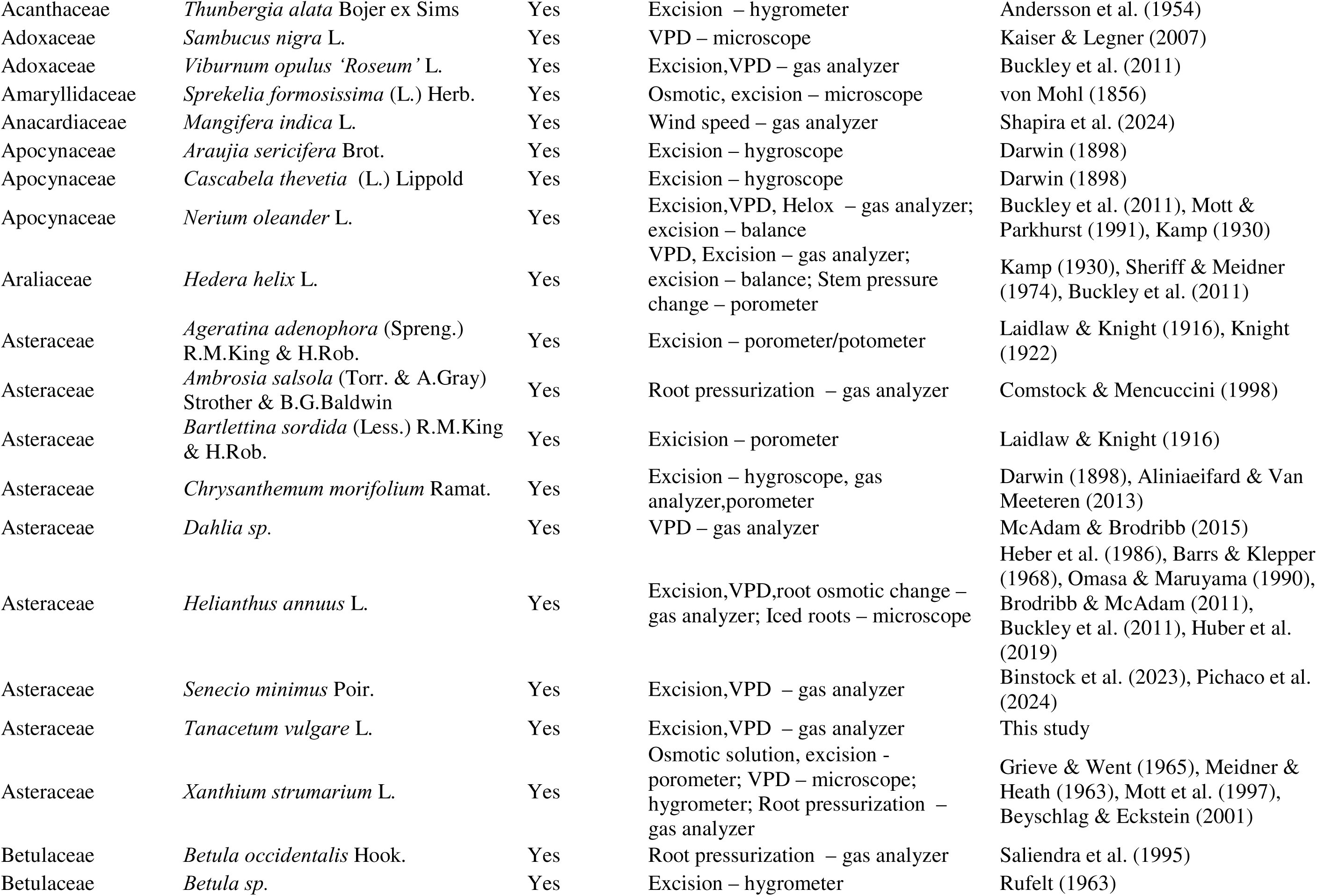
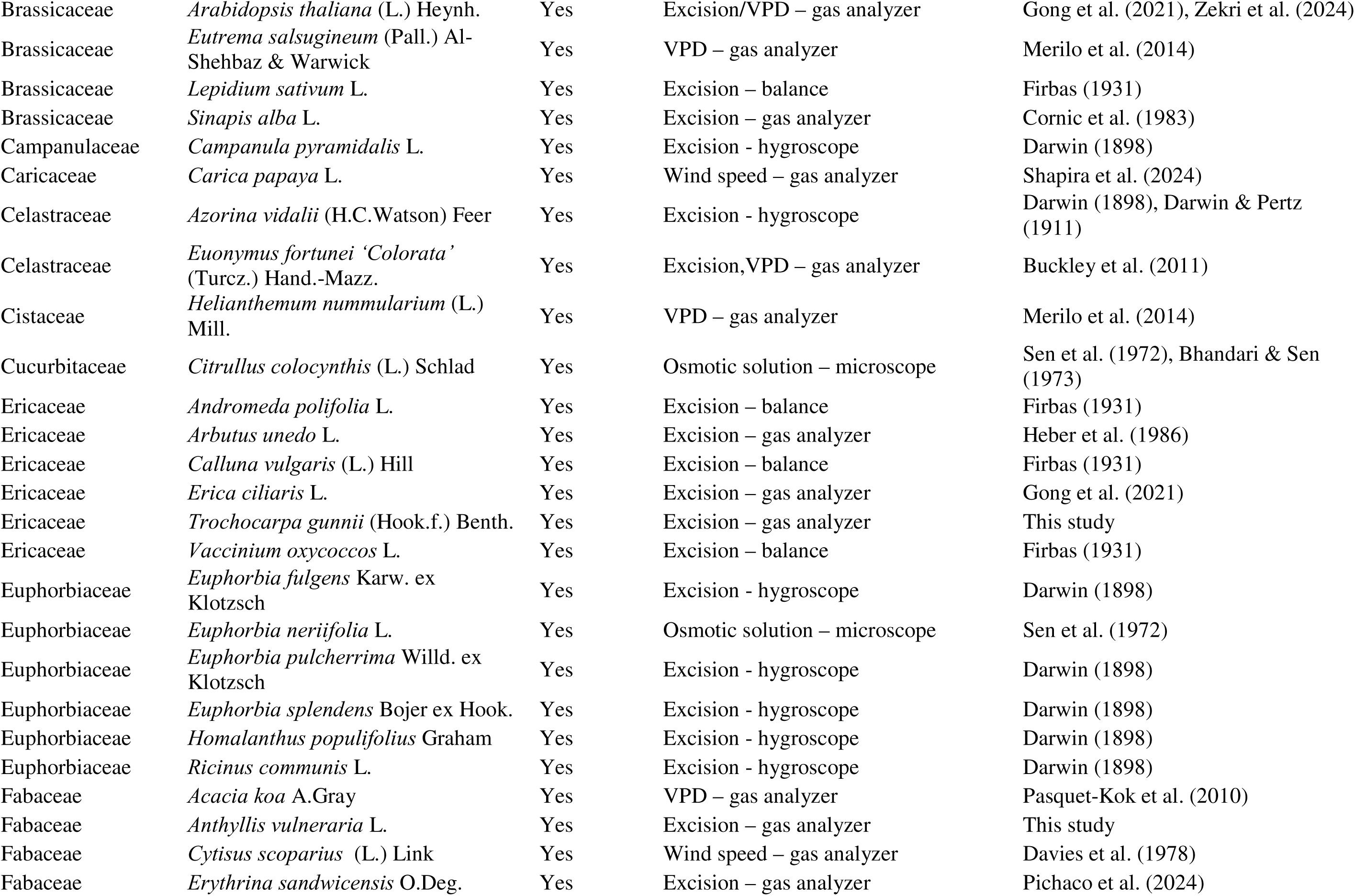
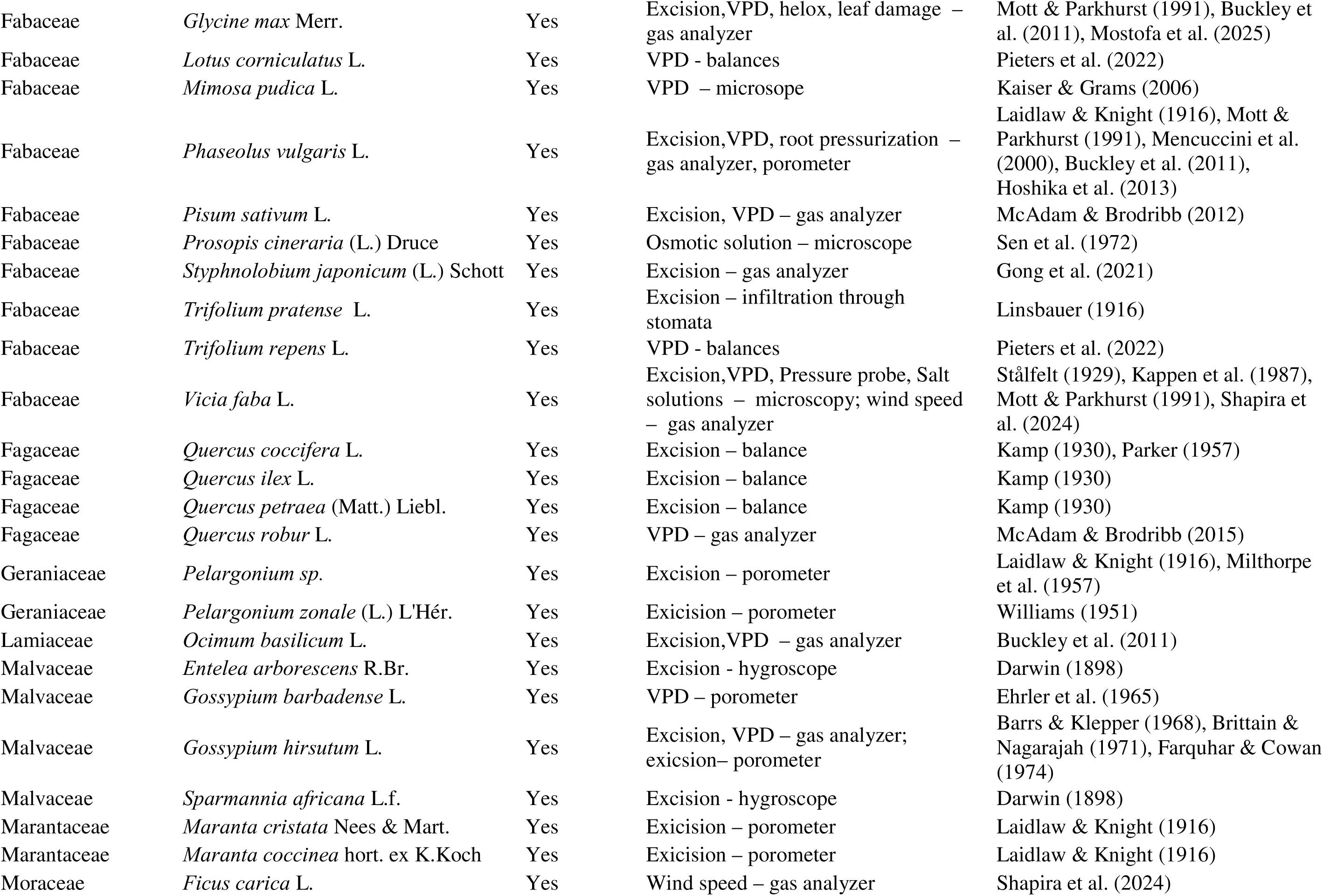
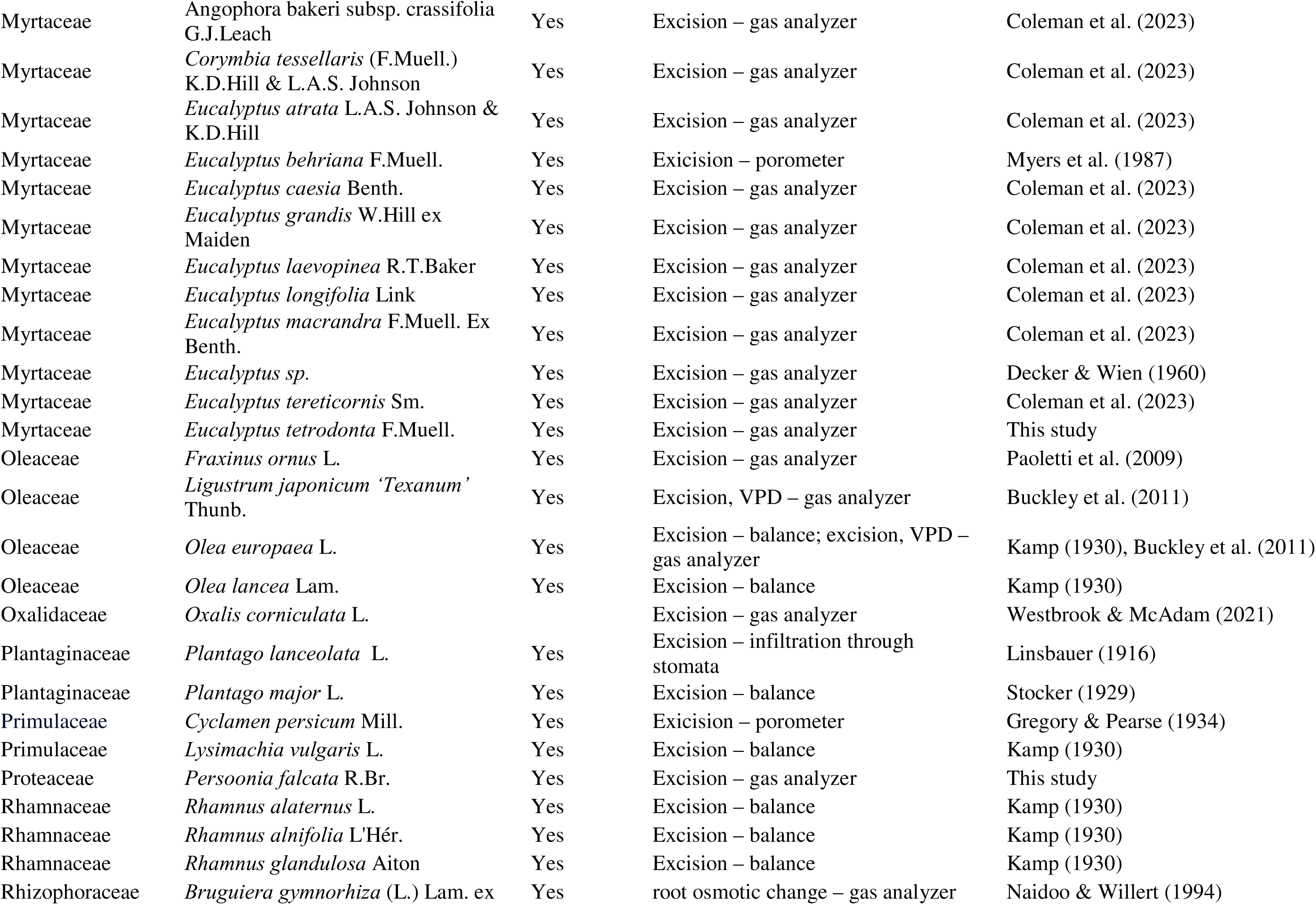
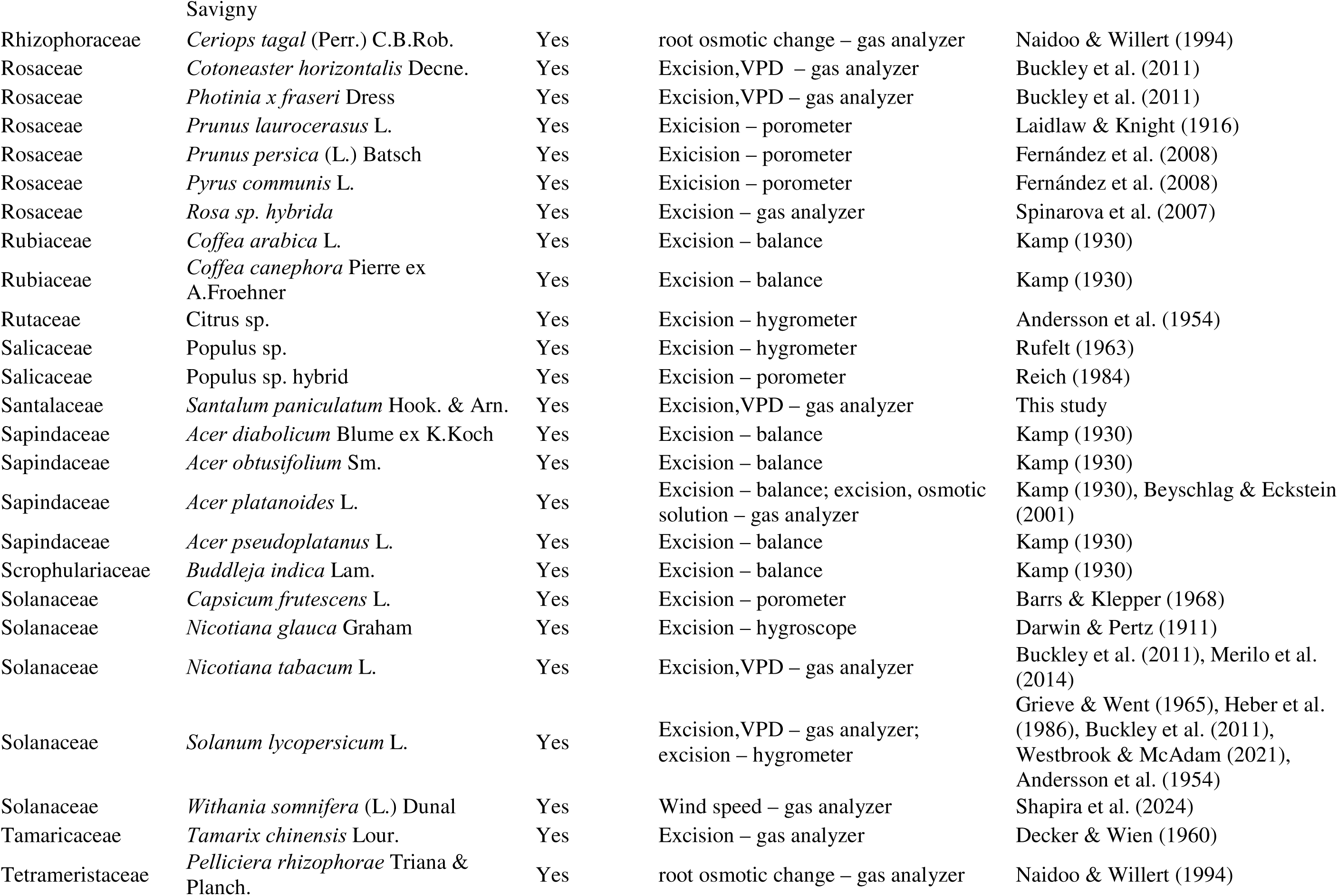
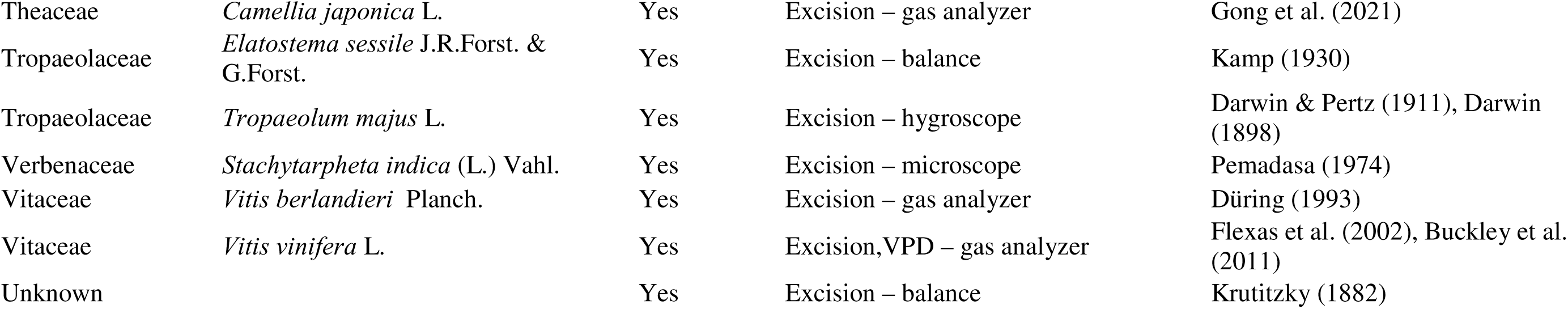
Species list of plants with or without mechanical advantage. Presence of mechanical advantage was determined based on the response of stomatal conductance or stomatal aperture to leaf excision, step increases in VPD, changes in epidermal or guard cell turgor pressure in leaf discs or epidermal peels, root pressurization induced increased whole plant hydraulic conductance or differences in guard cell and adjacent epidermal sizes in Cryo-SEM images.

### Plant material

The presence of mechanical advantage was tested in *A. trichopoda* grown in an open-air shadehouse without environmental control at the University of Tasmania, Sandy Bay, Tasmania, Australia. Three species belonging to the order Austrobaileyales (*Austrobailyea scandens, Schisandra chinensis, Illicium parviflorum*), one species in the Nymphaeales (*Nymphaea thermarum*), and three Magnoliids (*Calycanthus chinensis, Liriodendron tulipifera* and *Asimina triloba*) were sampled from the living collections at the Arnold Arboretum, with the exception of*L. tulipifera* which grew in a research garden in Cambridge, MA, USA. Gymnosperm representatives included *Ginkgo biloba* which grew on the grounds of Harvard University, Cambridge, MA, USA; *Cycas armstrongii* sampled in the field at Gunn Point, Northern Territory, Australia; and a potted individual of *Agathis robusta* which grew in an open-air shadehouse at the University of Tasmania. The eudicot angiosperm species *P. falcata* and *Eucalyptus tetrodonta* were sampled in the field at Gunn Point; *Anthyllis vulneraria* and *Tanacetum vulgare* were sampled in the field at La Tour de Carol, France; *Santalum paniculata* was a potted individual grown in the greenhouses of Purdue University, West Lafayette, IN, USA; *T. gunnii* was sampled from a plant growing on the grounds of the University of Tasmania. The lycophyte species *Selaginella pallescens* was sampled in the field in Costa Rica. For the ANITA-grade species, an average of 15 leaves per species were measured with the exception of *A. trichopoda*, where only one sample was taken. For the lycophyte, gymnosperm species and eudicot angiosperm species one sample each was taken. Measurements on individuals growing outdoors were made in the summer in the Northern Hemisphere, the wet season in the tropics, or the spring in the temperate Southern Hemisphere, on newest fully expanded leaves or shoots.

### Stomatal response to leaf excision

Stomatal conductance (g_s_) was measured using Licor (Li-Cor, Lincoln NE, USA) gas exchange systems (6400,6400XT or 6800). Conditions inside the cuvette were set to match ambient CO_2_ levels and PAR incident on the leaves, with the reference stream maintained at or above 70% relative humidity. Stomatal conductance of each leaf was monitored for at least three minutes to ensure stability (g_initial_, Supplementary Figure S3). The petiole was then cut in air to sever the water flow into the leaf and induce rapid dehydration. Measurements were recorded at 20 to 30 second intervals until stomatal conductance decreased to below 50% of the initial value.

A WWR was scored as present if leaf excision caused stomatal conductance to increase above the 95% confidence interval of the average g_initial_ during the three-minute period prior to leaf excision (Supplementary Figure S1). For leaves with a WWR, the maximum g_s_ post leaf excision (g_peak_) and the time after excision to reach g_peak_ (time to peak) was recorded. The WWR magnitude was calculated as the difference between g_peak_ and g_initial_. For measurements in which there was a WWR, the rate of change in g_s_ after leaf excision (mol m^-2^ s^-2^) was calculated from the linear regression between g_s_ and time between leaf excision and g_peak_. In the absence of a WWR the rate of g_s_ change over four minutes following leaf excision was calculated.

### Stomatal architecture

Stomatal density, index and size, as well as the arrangement of the surrounding epidermal cells was characterized for five leaves per species. We categorized epidermal arrangements as either paracytic or non-paracytic (Supplementary Figure S2). In a paracytic arrangement a pair of epidermal cells are arranged parallel to guard cells while all non-paracytic arrangements have two or more epidermal wall junctions adjacent to guard cells, potentially impeding the guard cell lateral movement when stomata open.

To prepare samples for imaging, leaves were fixed, preserved, cleared to remove pigmentation that would decrease the visibility of the boundaries between epidermal cells, and stained when the boundaries epidermal cells were indistinct. Leaf sections (0.5 cm^2^) were cut and stored in 7:1 ethanol and acetic acid fixative until samples could be imaged. For *N. thermarum*, *C. chinensis*, and *S. chinensis*, rinsing the stored samples with fixative solution three times with an interval of one hour between rinses was sufficient to clear the leaf segments. Leaf segments of *L. tulipifera, A. scandens*, *A. triloba* and some *S. chinensis* samples required additional steps to eliminate chlorophyll and other pigments. Leaf samples stored in ethanol and acetic acid solution were returned to aqueous solution using an ethanol dilution series (80%, 60%, 40%, 20%,0%) with ten minutes between each step. The samples where then put in 1M KOH aqueous solution overnight and rinsed with water. If the leaf segments still had color (e.g., some of the *A. triloba* samples) they were placed in 2% bleach for five minutes. *I. parviflorum* leaf segments were cleared using benzyl benzoate because the cuticle obscured the boundaries between epidermal cells even after clearing with the KOH protocol. Leaf segments stored in ethanol and acetic acid solution were transferred sequentially between a 2:1 solution of 100% ethanol to benzyl benzoate, a 1:1 solution of 100% ethanol to benzyl benzoate, and a 1:1 solution of benzyl benzoate to dibutyl phalate, with an hour interval between solution changes. After clearing, each sample was rinsed three times with water, with five minutes between rinses, and allowed to rest in water for an hour before mounting on a glass slide with glycerol. The cleared *A. triloba* leaves were stained for five seconds in 1% toluidine blue because the boundaries between epidermal cells were only faintly visible in the cleared leaf segments. For *A. trichopoda* leaf segments were treated in 2% bleach overnight until the epidermal layers could be isolated. The epidermal peels were then stored in water before imaging.

The abaxial epidermis of each segment was imaged using a Zeiss AxioImager; when the epidermal plane was too uneven to get a clear image of the leaf surface, an LSM700 Confocal microscope was used. The length of eight stomata per leaf was measured and stomatal density (stomata per unit area) and stomatal index (stomata per number of epidermal cells) determined from three 0.4 mm^2^ images taken from regions between the major veins. All of the stomatal complexes for each image were classified as paracytic or non-paracytic according to the arrangement of surrounding epidermal cells (Baranova & Baranova, 1992). A stomatal complex was considered paracytic if there were only two neighboring epidermal cells aligned laterally with the guard cells (Supplementary Figure S4).

## Results

A literature search revealed that WWRs have been observed in at least 173 species of angiosperm from 63 families (Figure 1, Table 1) based on reports from 201 publications beginning as early as 1856. Most of reports of the presence of mechanical advantage and WWRs were in the monocots and the eudicots. Outside of these major angiosperms groups there have been only two species in the Magnoliids that were reported to have WWRs, with no studies showing WWRs among the ANITA grade species until this study (Figure 1, Table 1).

**Figure 1.**
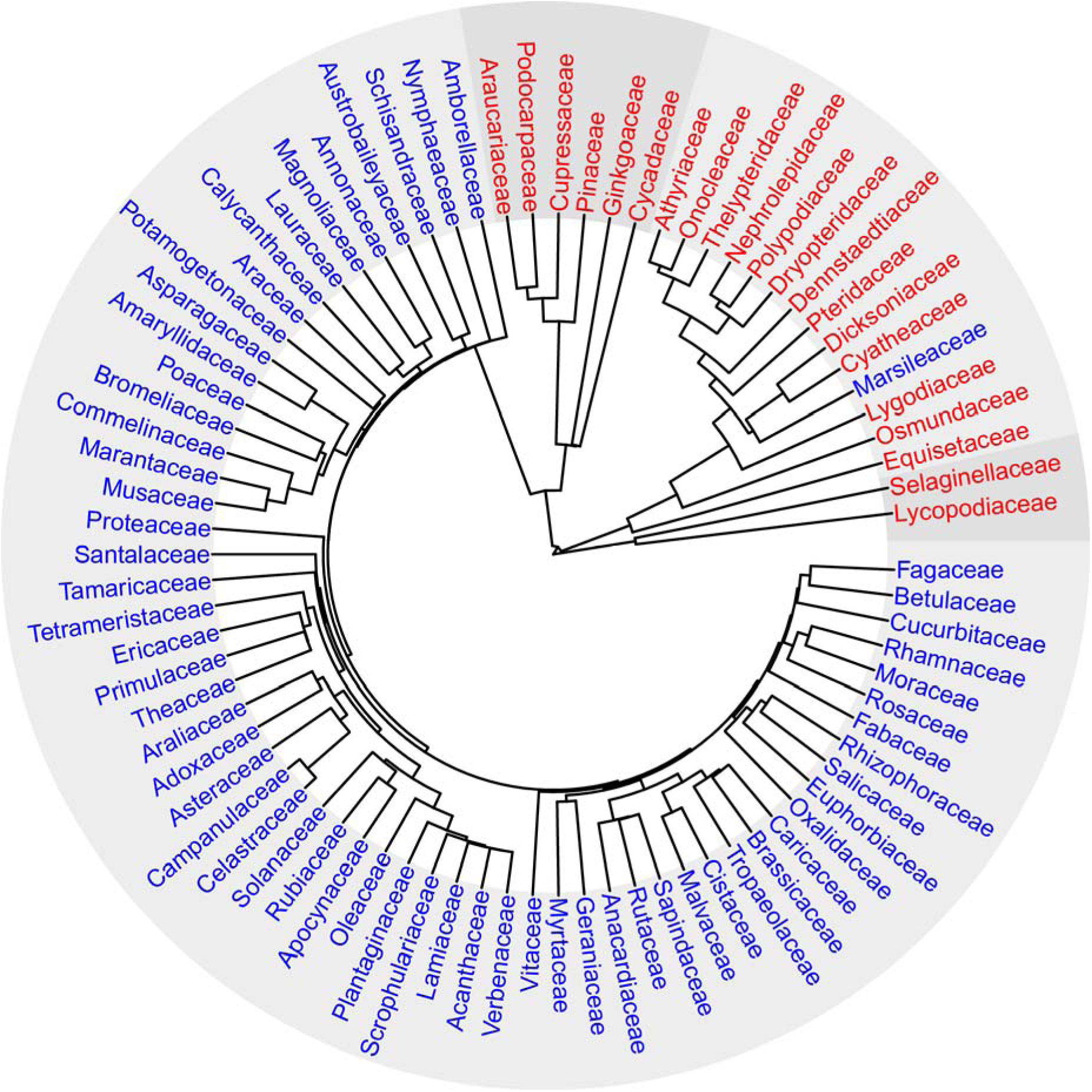
Phylogeny of vascular plant families where mechanical advantage has been tested. Blue indicates the presence of mechanical advantage and red an absence of mechanical advantage. The transitions between dark and light gray shading indicates a separation of groups from lycophytes, ferns, gymnosperms and angiosperms.

In this study, a transient increase in stomatal conductance, g_s,_ after leaf excision was observed in all ANITA grade species (Figure 2). Similar wrong-way stomatal opening was also observed in the three Magnoliid species and six eudicot species tested (Figure 2). Among the measured ANITA grade species in this study, WWR magnitudes sometimes exceeded 50% of g_initial_, the g_s_ before excision, as seen in *A. scandens* (Table S1), similar to the magnitudes in the eudicot *P. falcata* and magnoliid *A. triloba* (Figure 2). The WWR was observed soon after leaf excision and lasted around 2-11 minutes (time to g_peak_, Table S1) across angiosperm species. WWRs were absent in the three gymnosperm species measured, with an absence of WWR now reported in 15 gymnosperm species from six families (Figure 1, Table 1). With the exception of five species from two genera of Marsileaceae, an absence of WWRs has been reported so far in at least 43 fern and lycophyte species from 16 families (Figure 1, Table 1).

**Figure 2.**
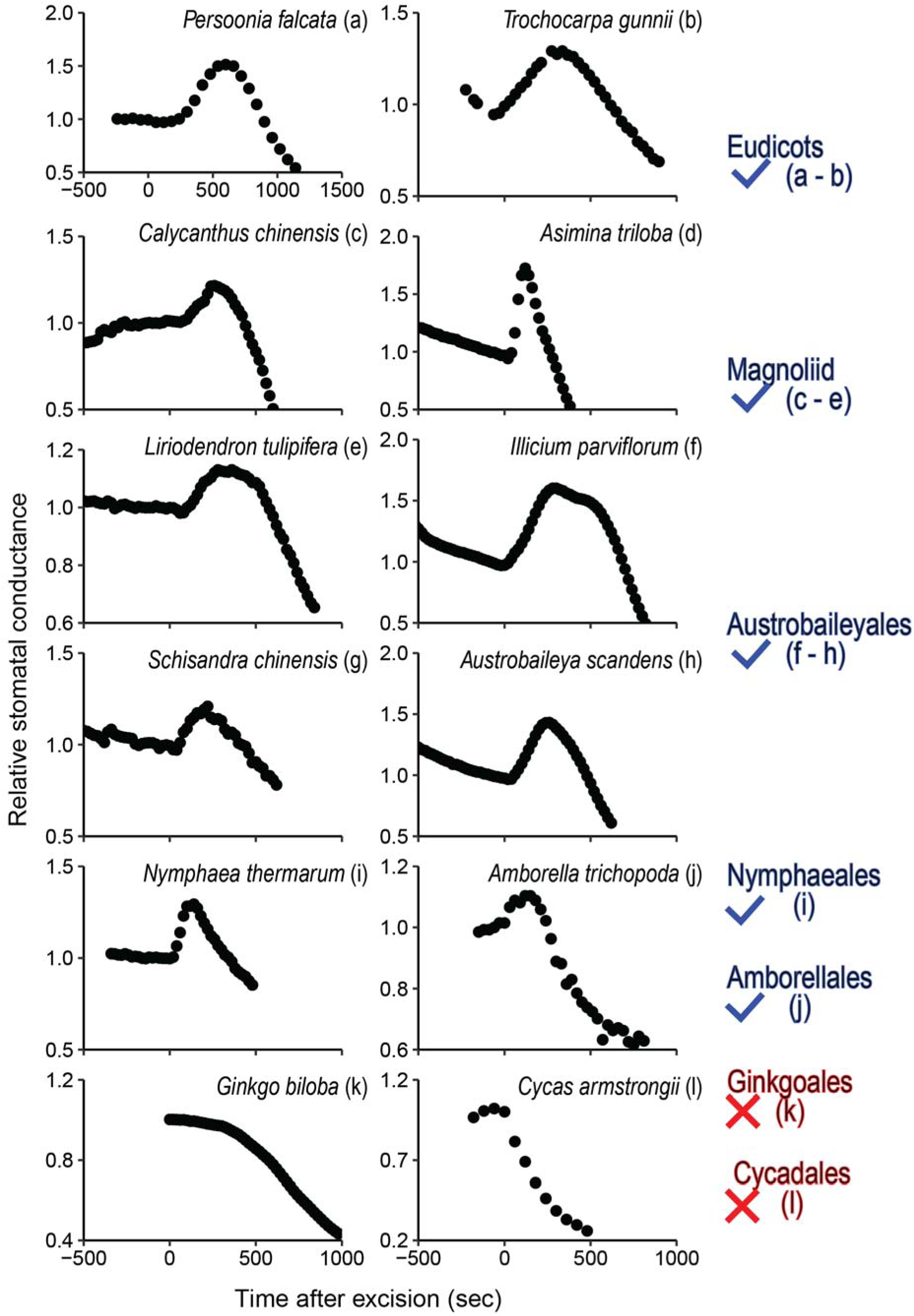
Representative relative stomatal conductance during leaf excision (at time 0) for each species along with the current distribution of wrong way response WWR and mechanical advantage in vascular plants. The cross marks a reported absence of WWR from species from that lineage taken from this figure or Table 1 and the tick marks the presence of WWR.

In angiosperms the magnitude of WWR is not a predictable trait since it varied leaf to leaf even for leaves sampled on the same day from the same individual (Figure 3). In some instances, leaf excision did not result in WWR (e.g., *L. tulipifera*, Figure 3, Supplementary Figure S3). There was variation in the frequency with which a WWR was observed across species (Table 2). In five ANITA grade species, a WWR occurred in more than 70% of the leaves sampled; 64% of the leaves sampled in *L. tulipifera* had a WWR and only 55% of the *N. thermarum* leaves had a WWR. Significant correlations in at least one of the measured species were observed when the initial slope of the g_s_ increase (mol m^-2^ s^-2^, Supplementary Figure S4) and the magnitude of the WWR (Supplementary Figure S5) were related to g_initial_, light intensity, and VPD, with the correlation consistently positive only for light intensity. In *L. tulipifera*, high VPD was negatively correlated with both the slope and the amplitude of the WWR.

**Figure 3.**
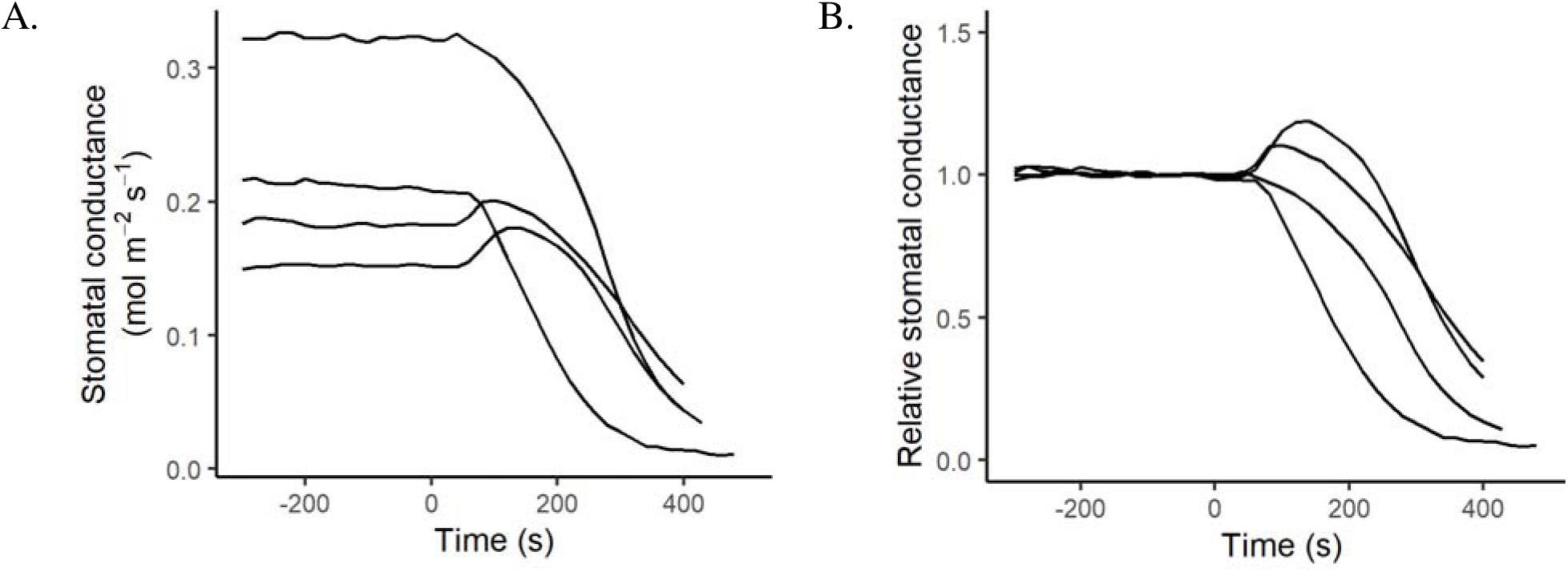
Variation in wrong way response (WWR) observations on leaves of the same individual of *Liriodendron tulipifera* measured on a single day. The traces include stomatal conductance before and after excision (A) and stomatal conductance expressed relative to the average stomatal conductance during the three minutes prior to leaf excision (B).

**Table 2.** Percentage of instances when WWR response was observed along with the percent of stomata with paracytic architecture.

| Species | Percent<br>positive<br>WWR (n) | Percent<br>Paracytic<br>complex |
| --- | --- | --- |
| *A. trichopoda* | 100 (1) | 82.9 |
| *A. scandens* | 95 (19) | 20.0 ± 5.0 |
| *S. chinensis* | 81 (21) | 32.5 ± 7.5 |
| *I. parviflorum* | 70 (10) | 97.5 ± 2.5 |
| *N. thermarum* | 55 (9) | 0.00 ± 0 |
| *L. tulipifera* | 64 (31) | 97.5 ± 2.5 |
| *A. triloba* | 75 (8) | 100 ± 0 |
| *C. chinensis* | 78 (9) | 97.8 ± 2.2 |

Often, a single leaf had a distribution of both paracytic and non-paracytic epidermal arrangement (Supplementary Figure S6). The percent of stomata with paracytic epidermal arrangement varied widely across the ANITA grade species even though all species have mechanical advantage (Table 2). There was no apparent relationship between stomatal architecture or size and the frequency of WWR exhibited by these species. Other stomatal characteristics varied markedly across the seven species. The three Austrobaileyales species had large (> 50 μm) and infrequent (< 60 mm^-2^) stomata, while the stomata of the three Magnoliids were smaller (< 32 μm) and more frequent (> 146 mm^-2^)(Supplementary Table S2). *N. thermarum* stomata were similar in size and density compared to the three Magnolids. The percentage of stomata with paracytic epidermal arrangement ranged from 100% in *A. triloba* to 0% in *N. thermarum*, with no apparent relation with taxonomic status, stomatal size, stomatal density or percent WWR observed.

## Discussion

Our observations and the results of the past 169 years of stomatal research find that mechanical advantage has evolved twice, once in the ancestor of angiosperms and once in the Marsileaceae. Mechanical advantage has been hypothesized to be a key mechanism in the achievement of high rates of gas exchange (Rockwell & Holbrook, 2017; Westbrook & McAdam, 2020). Since ANITA grade angiosperms largely have low stomatal conductance compared to other angiosperms (Feild *et al*., 2003) but also have mechanical advantage, the functional importance of mechanical advantage in these plants might instead allow faster stomatal opening in response to light flecks that occur in forest understories(Rockwell & Holbrook, 2017), as has been observed in species of Piperaceae (Chazdon & Pearcy, 1991; Tinoco-Ojanguren & Pearcy, 1992). A hydropassive WWR in response to a sudden increase in light intensity that increases the thermal load and leaf VPD could accelerate CO_2_ uptake (Delwiche & Cooke, 1977; Rockwell & Holbrook, 2017). Such opportunistic behavior is consistent with the hypothesis that angiosperms evolved in dark and disturbed environments (Feild *et al*., 2004). Thus, we hypothesize that the functional role of mechanical advantage could be twofold: wider stomatal opening and rapid higher sensitivity to environmental change like light (Lawson & Blatt, 2014; Rockwell & Holbrook, 2017).

A cost of WWRs is that stomatal responses are transiently disconnected from water status meaning that plant with mechanical advantage they require an active means of inducing stomatal closure in response to water stress (Westbrook & McAdam, 2021). Since stomatal closure in *A. trichopoda* at high VPD is mediated by ABA (McAdam & Brodribb, 2015), this mechanism is the likely safety valve for excessive stomatal opening from WWR when the leaf is challenged by conditions in which the demand for water to exceeds hydraulic supply.

WWR appear to be absent in gymnosperms (Rufelt, 1963; Mcadam & Brodribb, 2012; McAdam & Brodribb, 2014, 2015) (Figure 2). Gymnosperms also do not use ABA to initiate stomatal closure in response to rapid increases in VPD (McAdam & Brodribb, 2015; Sussmilch *et al*., 2017), consistent with an absence of mechanical advantage. The only occurrence of mechanical advantage outside the angiosperms is in the fern family Marsileaceae, which lack stomatal responses to ABA. In Marsileaceae excision causes stomata to open, persisting until epidermal turgor loss removes the effect of mechanical advantage and permits stomata to passively close (Westbrook & McAdam, 2021). The presence of mechanical advantage in Marsileaceae is likely a convergence with angiosperms that allows large stomatal conductances and high productivity. Being aquatic or semi-aquatic, the Marsileaceae never experience conditions that require complete stomatal closure. The lack of ABA-mediated stomatal closure likely prevents the spread of these ferns beyond the semi-aquatic habitats they are found in. Terrestrial plants would face catastrophic water loss if they had mechanical advantage without an active mechanism for stomatal closure as demonstrated by the excessive VPD-caused desiccation of ABA-deficient tomato mutants (Brodribb *et al*., 2021).

In angiosperms the substantial within and between species variation in the magnitude and especially in the consistent presence of the WWR could not be attributed to leaf epidermal architecture or plant-to-plant differences. Buckley et al. (2011) report wide variation in WWR among and within species and attribute this to differences in mechanical advantage, while Powles et al. (Powles *et al*., 2006) note that variation in WWR durations could be caused by delayed initiation of active stomatal closure mechanisms. WWR in angiosperms can be highly sensitive to VPD conditions and windspeed (Zait *et al*., 2017; Pichaco *et al*., 2024; Shapira *et al*., 2024) leading to variation in transient opening speeds and magnitudes. The measurements made in this study were under natural conditions such that the within-species variability seen in this study is to be expected.

The anatomical basis for the presence of mechanical advantage does not lie in epidermal arrangement around the guard cells. We find that the frequency of paracytic arrangements does not influence mechanical interactions and that the percent of WWR response observed was not related to the percent of paracytic arrangements (Table 2). The high degree of epidermal cell wall sinuosity and hence epidermal cell wall rigidity in *T. gunnii* also did not impede WWR (Figure 2). Though specialization of paracytic subsidiary cells in the grasses is likely important for stomatal opening (Lawson & Vialet□Chabrand, 2019), epidermal arrangement cannot be a diagnostic tool for the presence of mechanical advantage. In species with mechanical advantage, epidermal cell size but not arrangement, influences the speed of stomatal opening under high VPD conditions (Pichaco *et al*., 2024). The search for the epidermal traits that allow mechanical advantage is ongoing and needs further research. One trait being the ratio of guard cell and epidermal cell lumen depth along their shared wall, which has been used for a geometric interpretation of mechanical advantage(Wu *et al*., 1985). Another promising epidermal trait that must be considered is the cellulose orientation in guard cell walls since regions of high wall stiffness places ferns, angiosperms with allantoid guard cells, and monocots with dumbbell guard cells into three distinct groups (Shtein *et al*., 2017).

Mechanical advantage is present in ANITA grade angiosperms and is widespread among angiosperms, but not in other groups of vascular land plant. Though among angiosperms, the ANITA grade species have comparatively low maximum stomatal conductance, the presence of mechanical advantage might allow plants to optimize gas exchange in response to increases in VPD due to light. Though epidermal arrangements around the guard cells are diverse across angiosperms, the presence or absence of distinct lateral epidermal cells bracketing the guard cells or epidermal cell sinuosity does not appear to be connected to the presence of mechanical advantage or the consistency or magnitude of WWR.

## Acknowledgements

We thank Greg Jordan for his recommendation for the sampling of *Trochocarpa*; Andrey Ostrovsky and Ms Bozhkova from the St. Petersburg Society of Naturalists for providing translation and information on the oral presentation by Krutitzky; David Bowman for providing logistical support for the sampling of species at Gunn Point; the Arnold Arboretum at Harvard for allowing us to sample from the living collection. This work was supported by Australian Research Council grants to TJB and SAMM, as well as National Science Foundation grants (IOS 2333890, IOS 23333888 and DMR 2011754) to SAMM and NMH.

## Author contributions

AM, NH, FR and SM designed different aspects of the research. AM and SM conducted the literature review and the experiments with the help of YF to collect epidermal anatomy data. AM analyzed the data and wrote and edited the manuscript with the help of SM, NH, FR and TB.

## Data availability

All data is available upon reasonable request.

