## Supplemental information for "The origin of mechanical advantage in angiosperms"

### Supplementary tables and figures:

Table S1: The measured initial stable stomatal conductance g_initial_, the magnitude of wrong way response (WWR) as a percentage of g_initial_, the time it took for the WWR to reach the maximum stomatal conductance, g_s_ and the photosynthesis levels measured during g_initial_. Averages and the standard error is presented when sample availability allowed more than one measurement. Averages are of measurements made on six leaves from two to five individual plants.

| Species | g_initial_ (mol m^-2^ s^-1^) | Percent increase in g_s_ (%) | Time to g_peak_  (minutes) | A_stable_  (µmol m^-2^ s^-1^) |
| --- | --- | --- | --- | --- |
| *A. trichopoda* | 0.137 | 8.8 | 2.0 | 6.92 |
| *A. scandens* | 0.051 ± 0.004 | 67 ± 4.8 | 11.2 ± 1.4 | 3.31 ± 0.07 |
| *S. chinensis* | 0.088 ± 0.008 | 28 ± 1.9 | 5.00 ± 0.55 | 6.79 ± 0.12 |
| *I. parviflorum* | 0.108 ± 0.025 | 22 ± 4.9 | 5.04 ± 0.89 | 5.14 ± 0.09 |
| *N. thermarum* | 0.466 ± 0.067 | 10 ± 2.1 | 2.80 ± 0.36 | 5.16 ± 0.26 |
| *L. tulipifera* | 0.184 ± 0.027 | 57 ± 2.9 | 3.07 ± 0.37 | 13.40 ± 0.18 |
| *A. triloba* | 0.053 ± 0.007 | 45 ± 3.4 | 1.71 ± 0.122 | 6.95 ± 0.12 |
| *C. chinensis* | 0.103 ± 0.015 | 13 ± 1.2 | 3.24 ± 0.82 | 9.09 ± 0.46 |

Table S2. A summary of stomatal complex characteristics for each species. The number of stomatal complexes without epidermal radial walls in contact with the ventral walls of the guard cells are represented as % paracytic complex.

| Species | Stomatal length (μm) | Stomatal index  (%) | Stomatal density  (mm^-2^) |
| --- | --- | --- | --- |
| *A. trichopoda* | 30.9 ± 0.22 | 17.6 | 173 ± 4.6 |
| *A. scandens* | 50.6 ± 0.56 | 9.6 | 58.8 ± 1.7 |
| *S. chinensis* | 55.2 ± 0.64 | 9.0 | 54.9 ± 2.7 |
| *I. parviflorum* | 53.7 ± 0.66 | 7.3 | 58.4 ± 1.8 |
| *N. thermarum* | 25.2 ± 0.74 | 9.8 | 151 ± 9.8 |
| *L. tulipifera* | 31.8 ± 0.62 | 15.7 | 146 ± 7.6 |
| *A. triloba* | 28.7 ± 0.34 | 14.6 | 298 ± 6.3 |
| *C. chinensis* | 24.8 ± 0.38 | 14.6 | 304 ± 19 |

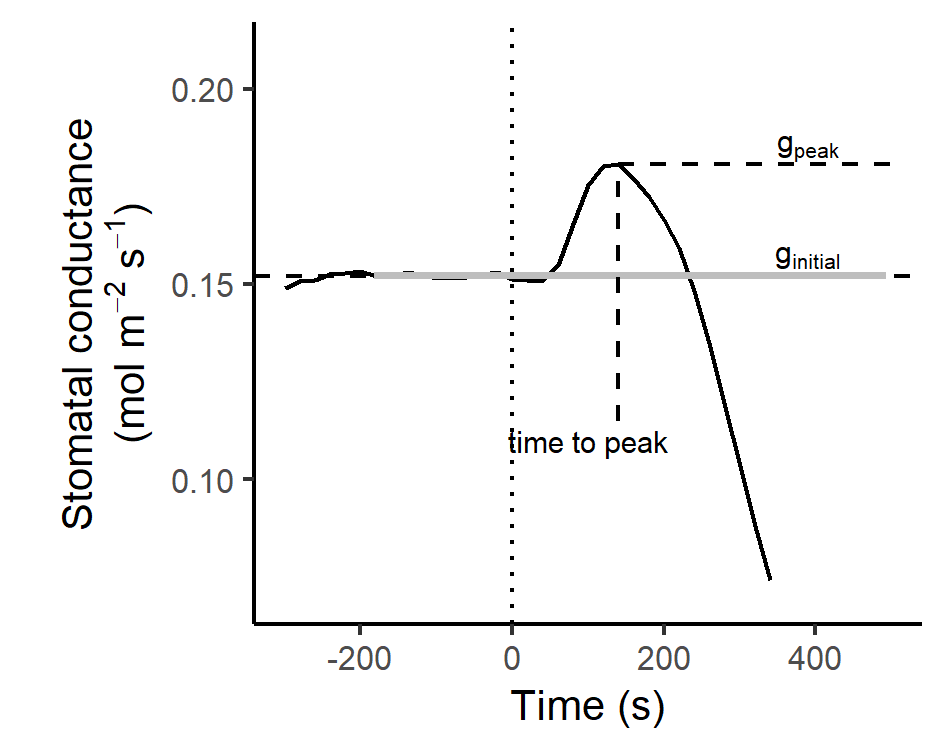

Figure S1. A representative trace of stomatal conductance over time illustrating a wrong way response to petiole excision in air. The dotted line at time zero is the point of leaf excision; g_initial_ is the average stomatal conductance over a three-minute duration before the leaf was excised. The gray bar on the dashed line designating g_initial_ represents the 95% confidence interval around g_initial_.

| 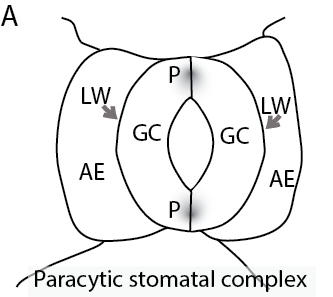 | 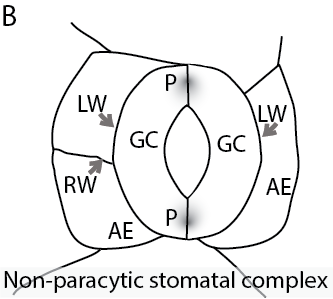 |
| --- | --- |
| Figure S2. Diagram of a paracytic (A) and a non-paracytic (B) stomatal complex, with guard cell (GC) polar regions (P), lateral walls (LW) and radial walls (RW) of adjacent epidermal (AE) cells, epidermal cells that are in contact with the guard cells Stomatal complexes with guard cells only in contact with the lateral walls of two adjacent epidermal cells are considered paracytic and if there are any radial walls in contact with the non-polar regions of the guard cells then the stomatal complex is not paracytic. | |

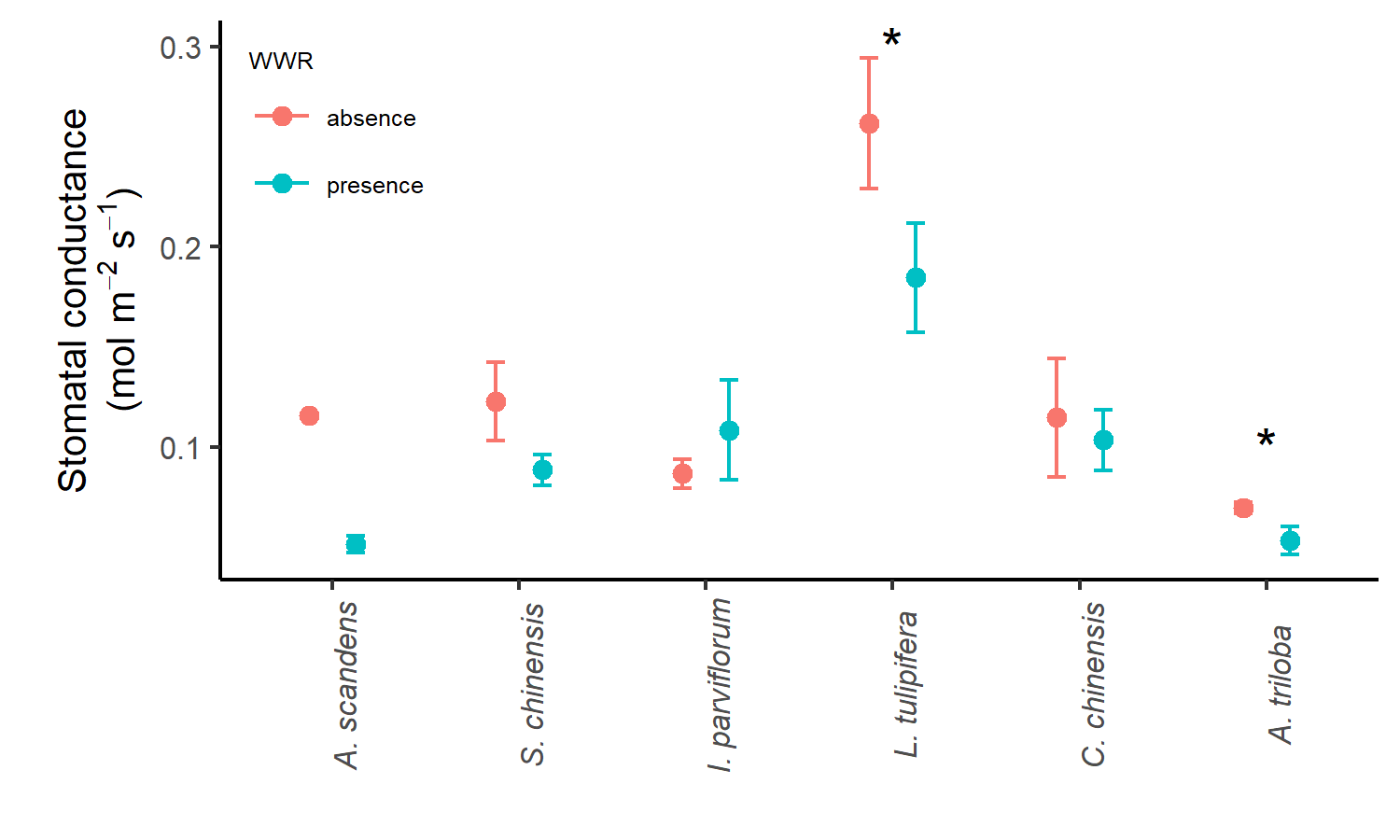

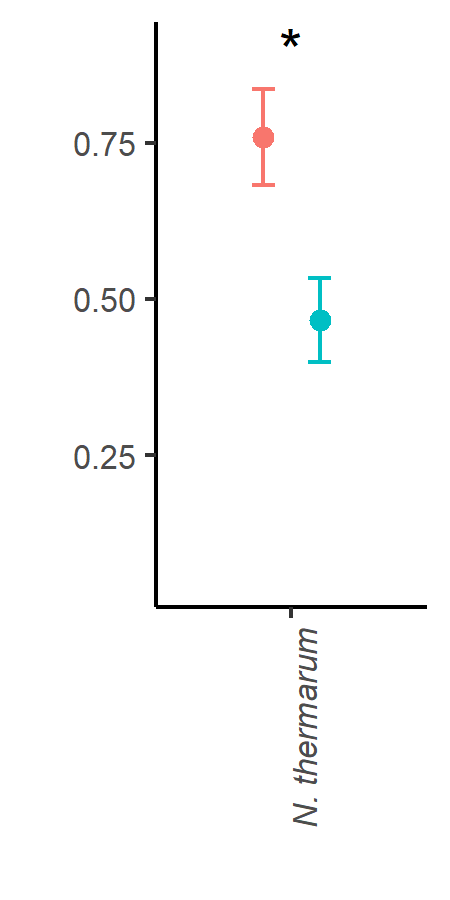

Figure S3. The average stomatal conductance (g_initial_) before leaf excision for each species group by the presence or absence of wrong way response. The bars represent the standard error for each species mean. The ‘*’ denotes differences in mean g_initial_ between leaves with and without WWR (p < 0.05 for the Welch one-sided t-test). Note that *A. scandens* had only one leaf without a wrong way response. *N. thermarum* is plotted against a wider y-axis.

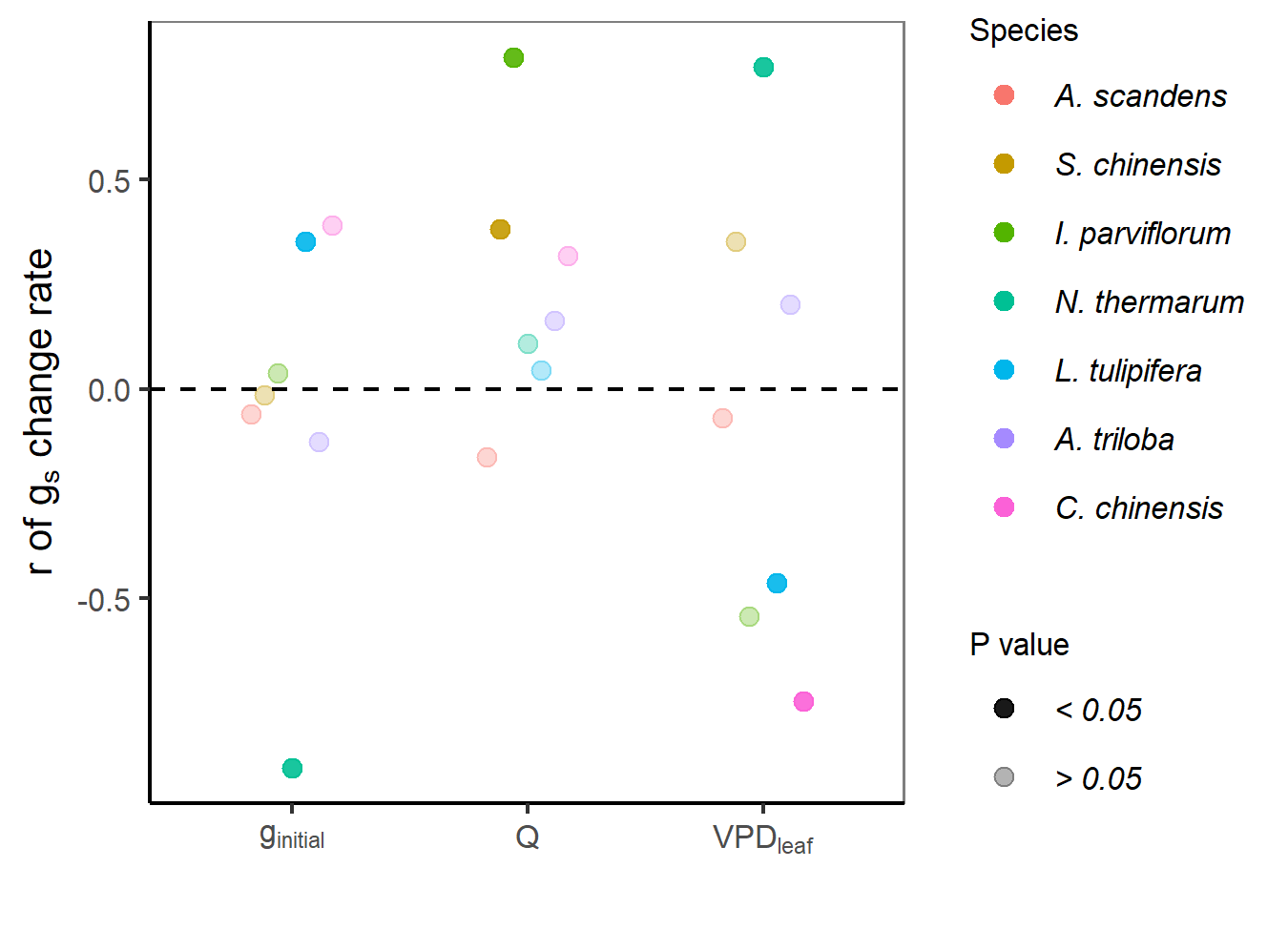

Figure S4. The correlation between the rate of change in stomatal conductance following leaf excision (mol m^-2^ s^-2^) and pre-excision stomatal conductance (g_initial_), light intensity (Q), and vapor pressure deficit (VPD_leaf_).

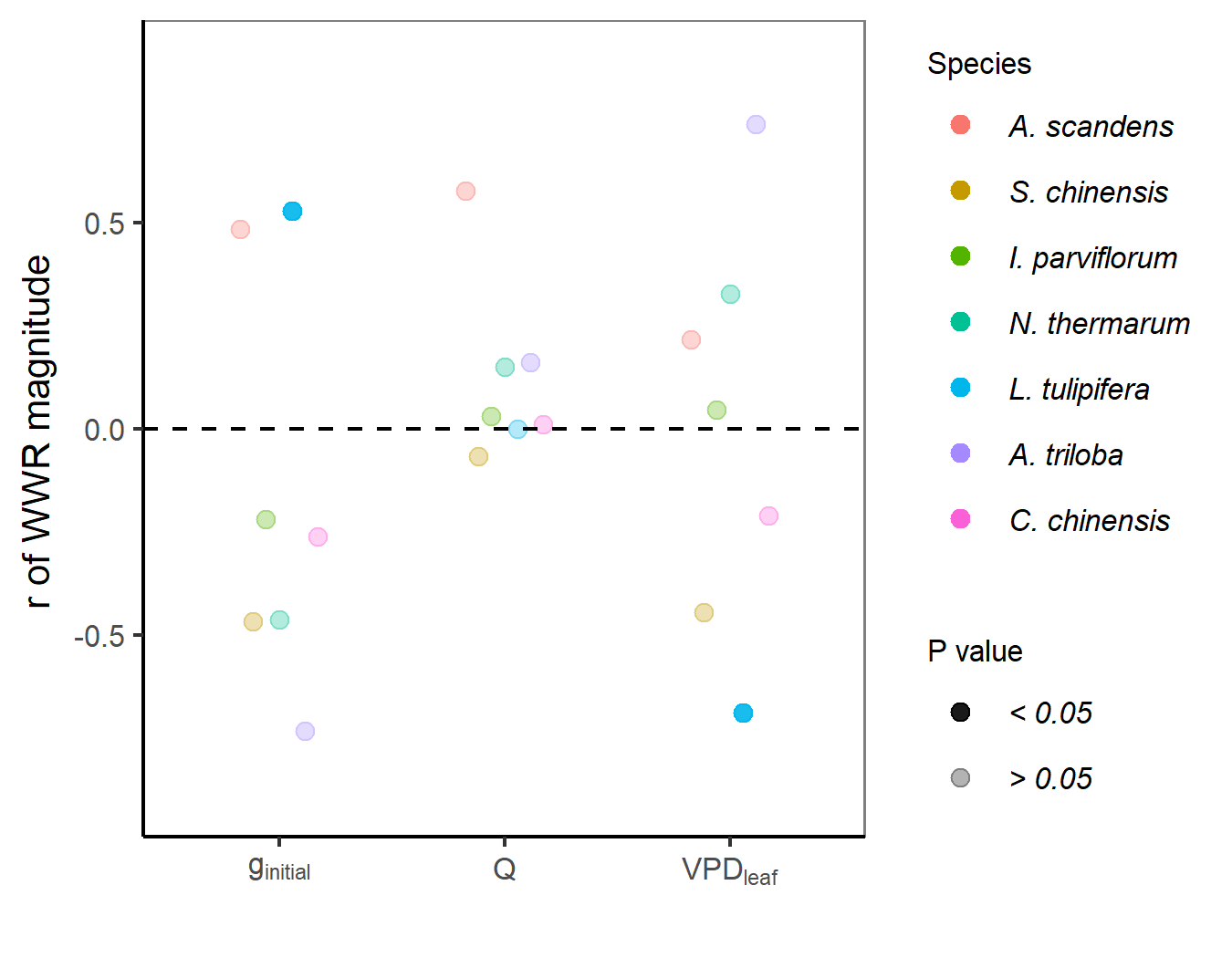

Figure S5. The correlation between initial stomatal conductance (g_initial_), VPD around the leaf measured and the magnitude of the WWR (g_peak_ - g_initial_).

| A. | 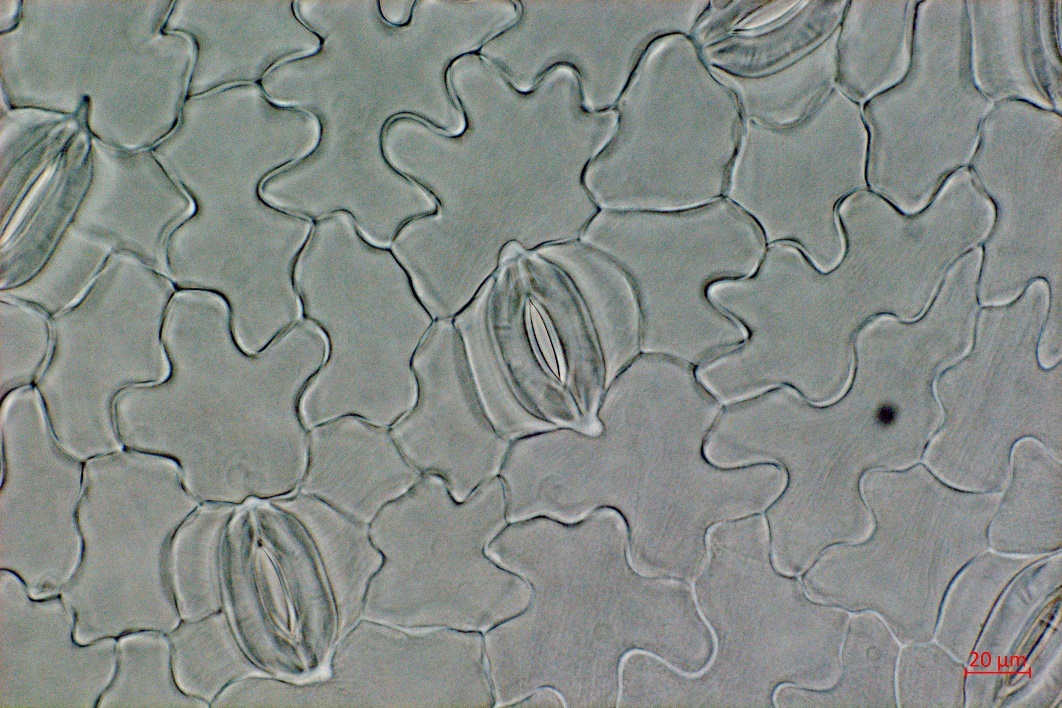 | B. | 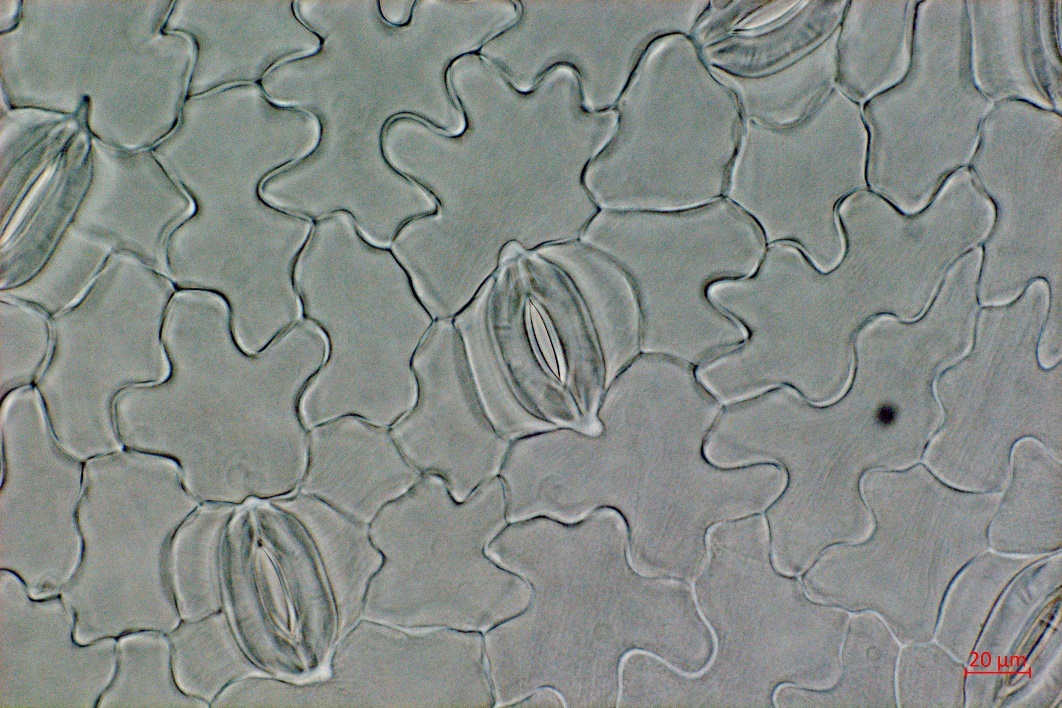 |
| --- | --- | --- | --- |
| Figure S6. (A) A paracytic stomatal complex and (B) a non-paracytic stomatal complex in S. chinensis. | | | |

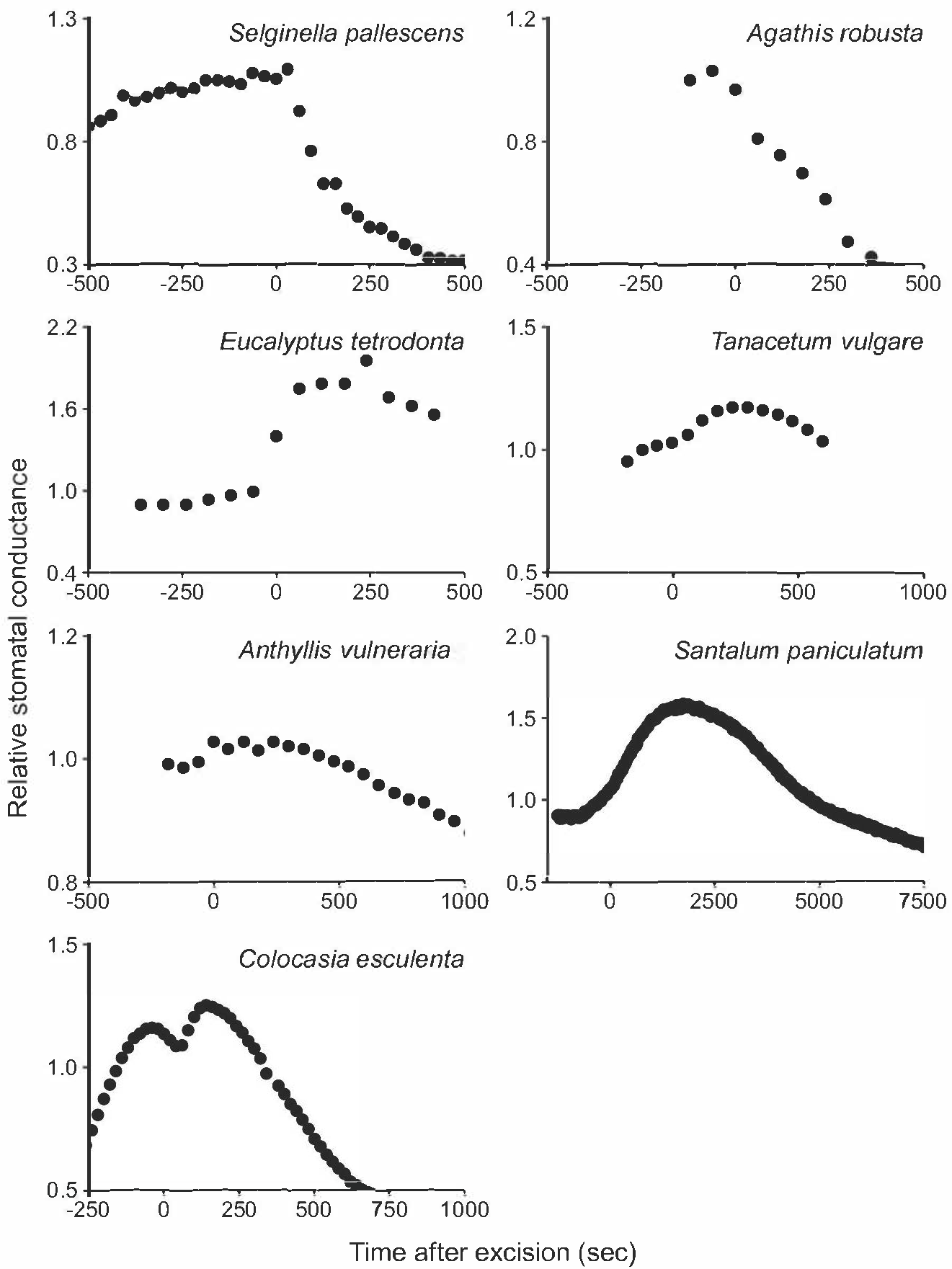

Figure S7. Leaf excision traces across species in which the presence of mechanical advantage had not been tested. The stomatal conductance is relative to the initial stomatal conductance before excision, g_initial_.
